# Single-cell transcriptomic changes in oligodendroglial lineage cells derived from Parkinson’s disease patient-iPSCs with LRRK2-G2019S mutation

**DOI:** 10.1101/2024.07.01.601392

**Authors:** Mohammad Dehestani, Wiebke Kessler, Nasser Karmali, Wenhua Sun, Polina Volos, Stanislav Tsitkov, Ashutosh Dhingra, Salvador Rodriguez-Nieto, Julia Tietz, David Schafflick, Noémia Fernandes, Julia Fitzgerald, Ernest Fraenkel, Thomas Gasser, Nisha Mohd Rafiq, Tanja Kuhlmann, Vikas Bansal

## Abstract

Despite extensive research, the contribution of the LRRK2 p.G2019S mutation to Parkinson’s disease (PD) remains unclear. Recent findings indicate oligodendrocytes (ODCs) and their progenitors are vulnerable in PD pathogenesis. Notably, oligodendrocyte precursor cells (OPCs) exhibit high endogenous expression of *LRRK2*. We induced PD patient-iPSCs with the LRRK2 p.G2019S mutation into oligodendroglial lineages and performed single-cell RNA sequencing. Cell type composition analysis revealed an increase in OPCs, proliferating OPCs and ciliated ependymal cells in LRRK2 lines, all of which are characterized by *LRRK2* expression. Differential expression analysis revealed transcriptomic changes in several pathways, including down-regulation of genes related to myelin assembly in ODCs, semaphorin-plexin pathway in OPCs, and cilium movement in proliferating OPCs. Cell-cell communication analysis identified significant alterations in several signaling pathways including a deactivation of PSAP signaling and an activation of MIF signaling in LRRK2 lines. Additionally, we observed an overall increase in SEMA6 signaling communication in LRRK2 cell lines; however, OPCs derived from these LRRK2 lines specifically lost SEMA6 signaling due to a down-regulation of *SEMA6A* and *PLXNA2*. Pseudotemporal trajectory analysis revealed that *SHH* had significantly altered expression along the pseudotime, accompanied by higher expression levels in LRRK2 lines. We propose that dysfunctional semaphorin-plexin signaling, along with cilia movement and SHH signaling, might represent early events in PD pathology.

## Introduction

Parkinson’s disease (PD) is the second most common age-related neurodegenerative disorder characterized by the loss of dopaminergic neurons in the substantia nigra region of the brain[1]. In fact, PD is the most common synucleinopathy to date that displays accumulation of aggregated form of α-synuclein protein encoded by *SNCA* gene. While the exact cause of PD remains elusive, genetic factors have been identified as significant contributors to the disease pathogenesis[2,3]. Among the various genetic factors associated with PD, the LRRK2 p.G2019S point mutation that results in abnormally high kinase activity, is recognized as one of the most prevalent variants worldwide implicated in the familial form of PD. In Ashkenazi Jewish and North African Arab populations, significantly higher prevalence was found, occurring in up to 20-40% of both familial and sporadic cases[3]. The LRRK2 p.G2019S mutation is a missense mutation in the leucine-rich repeat kinase 2 (*LRRK2*) gene, which encodes a multifunctional protein involved in various cellular processes, including mitochondrial homeostasis, autophagy, and lysosomal function[4,5]. Nearly two decades after linking *LRRK2* to PD[6,7], extensive research has been conducted to uncover the mechanistic pathways through which this mutation contributes to PD pathogenesis. The research particularly focuses on its effects on pathways such as mitophagy and lysosomal function in neurons, especially dopaminergic neurons[8,9].

Advanced high-throughput single-cell omics analyses have identified a significant association between PD genetic risk and oligodendroglial lineage cells[10–13] (including both oligodendrocyte precursor cells and oligodendrocytes), indicating their potential role in the development of the PD. Oligodendrocytes (ODCs), the myelinating cells of the central nervous system, are derived from oligodendrocyte precursor cells (OPCs), and play a crucial role in maintaining the integrity and function of axons, facilitating efficient signal transmission between neurons[14–16]. While traditionally recognized for their crucial role in myelination, ODCs and OPCs are now acknowledged for their wide and dynamic range of functions[16–20]. ODCs not only form dynamic myelin sheaths around axons, which play a crucial role in higher brain functions such as learning and memory[21,22] but also provide vital metabolic support to neurons[15], highlighting their contribution to overall brain function. OPCs are integrated into local neural circuits[23] and participate in synaptic[24] and axonal remodeling[25,26], phagocytosis process[27] and immunomodulation[28]. Intriguingly, it has been observed that the *LRRK2* gene is expressed at significantly higher levels in OPCs compared to other cell types in the substantia nigra (SN)[12,29,30]. Hence, a clear avenue of investigation would involve exploring the effects of the LRRK2 p.G2019S mutation on oligodendroglial lineage cells and their potential role in PD development and progression.

We aimed to identify molecular changes in oligodendroglial lineage cells derived from induced pluripotent stem cells (iPSCs) by comparing PD patients carrying the LRRK2 p.G2019S mutation with healthy controls (HC). By differentiating patient-specific iPSC lines to an oligodendroglial lineage, we were able to explore the genetic effects of the LRRK2 p.G2019S mutation at an unprecedented level and created a comprehensive molecular cell atlas using single-cell transcriptomics coupled with immunocytochemistry (ICC) (Figure 1A). Using state-of-the-art computational analysis methods, we found significant dysregulation of the genes related to pathways like chemotaxis, calcium-mediated signaling, cell-cell adhesion, myelin assembly, fatty acid transport, I-kappaB kinase/NF-kappaB signaling, mitochondrion disassembly, semaphorin-plexin, sonic hedgehog and cilia movement. We anticipate that these results will aid in understanding the disease mechanisms and identifying potential therapeutic targets in PD, with a particular focus on oligodendroglial cells.

**Figure 1:**
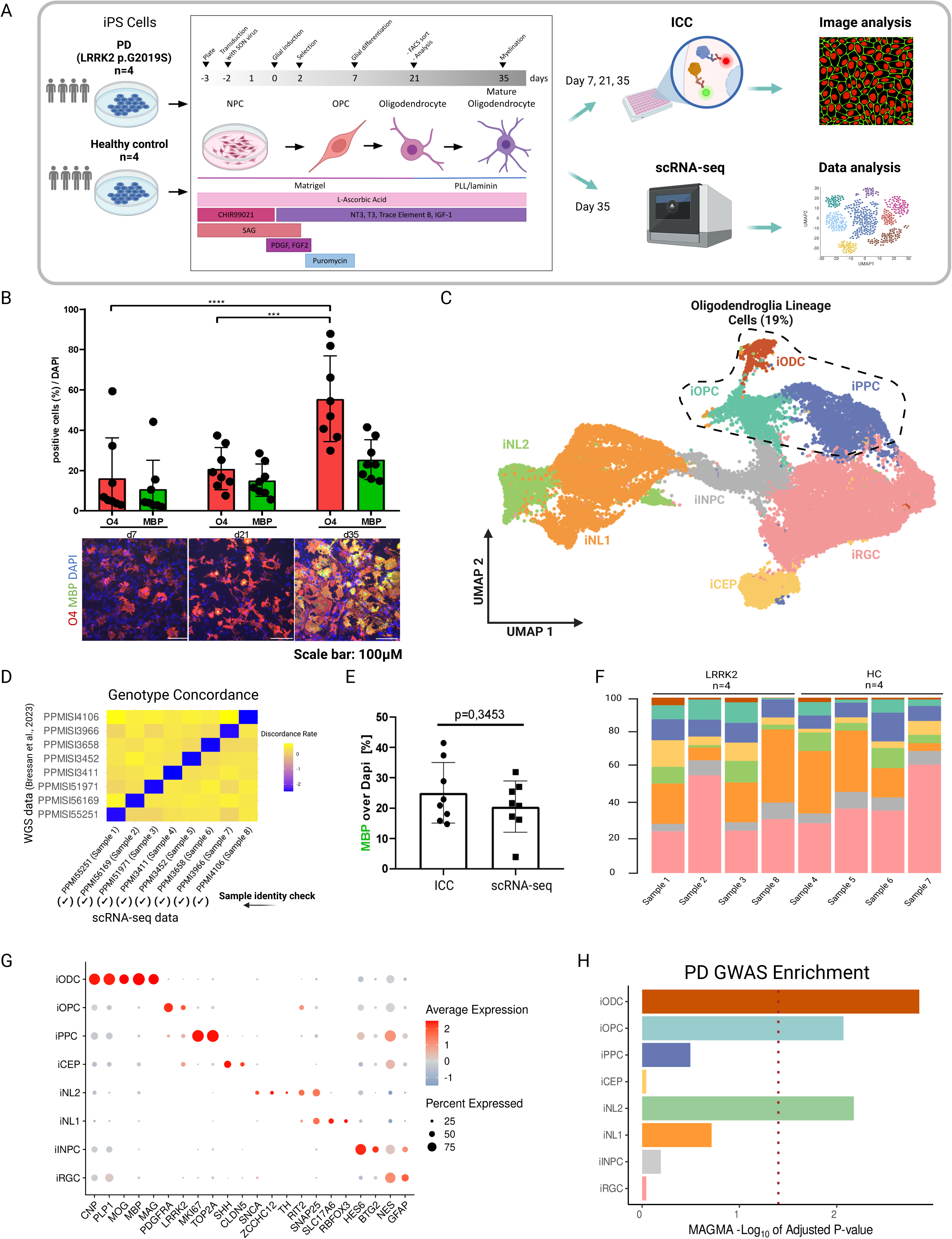
Overview of experimental plan, imaging and scRNA-seq analysis. **(A)** Schematic overview of the experimental plan. **(B)** Barplot representation of positive cells detected by ICC for oligodendrocyte markers O4 and MBP normalized to the total number of cells for day 7, day 21 and day 35 of the differentiation protocol. (**** represents p-value < 0.001 and *** represents p-value < 0.01, one-way ANOVA) **(C)** Uniform manifold approximation and projection (UMAP) visualization of the scRNA-seq clusters from 35,184 high quality cells. Cells are color-coded based on their assigned clusters; induced oligodendrocytes (iODCs), induced oligodendrocyte precursor cells (iOPCs), induced proliferative precursor cells (iPPCs), induced ciliated ependymal cells (iCEP), induced neuron-like cells 2 (iNL2), induced neuron-like cells 1 (iNL1), induced intermediate neuronal progenitor cells (iINPC) and induced radial glial cells (iRGCs). **(D)** Genotype concordance showing the consistency of genotypes between the scRNA-seq data and donor (blood-derived) whole-genome sequencing (WGS). **(E)** Percentage of MBP+ cells detected by ICC and scRNA-seq normalized to the total number of cells. Due to low *MBP* gene expression in sample 8, it was subsequently excluded from downstream analysis. **(F)** Barplot representing the distribution of cell-type percentages across sample and mutation. **(G)** Dotplot illustrating the expression of known gene markers for cell identity. **(H)** Multi-marker analysis of genomic annotation (MAGMA) gene set enrichment based on 29,559 high quality cells showed significant associations with induced-oligodendrocytes (iODCs), induced-oligodendrocyte precursor cells (iOPCs) and induced-Neuron-like 2 (iNL2) cells.

## Results

We used 8 induced pluripotent stem cells (iPSC) lines, comprising 4 healthy controls (HC) and 4 with the LRRK2 p.G2019S mutation, which will be referred to as Parkinson’s disease (PD) lines (Table S1). These lines were obtained from the Parkinson’s Progression Markers Initiative[31] (PPMI; https://www.ppmi-info.org/) and were also used in our previous consortium study, the Foundational Data Initiative for Parkinson Disease[32] (FOUNDIN-PD; https://www.foundinpd.org/). The iPSC lines were differentiated into oligodendroglial lineage cells using a previously published protocol[33], and their efficiency was evaluated through immunocytochemistry (ICC) at day 7, 21 and 35 (Figure 1A-B and Table S1). The O4^+^ cells (O4 antigen is a cell-surface sulfatide expressed on late-stage OPCs) were significantly increased from day 7 to day 35 and accounted for 55.6 +/- 7.5 percent of DAPI^+^ cells at day 35 (Figure 1B). Deviating results from previously published study could be due to the different cultivation conditions of the neuronal precursor cells (NPCs)[34–36]. Next, we conducted single-cell RNA sequencing (scRNA-seq) on differentiated iPSCs at day 35 (Table S1). Following data cleaning and quality control (see Methods), we analyzed 35,184 high-quality single cells (18,125 cells from PD and 17,059 cells from HC). Clustering analysis revealed 8 major cell clusters identified as induced oligodendrocytes (iODCs), induced oligodendrocyte precursor cells (iOPCs), induced proliferative precursor cells (iPPCs), induced ciliated ependymal cells (iCEP), induced neuron-like cells 2 (iNL2), induced neuron-like cells 1 (iNL1), induced intermediate neuronal progenitor cells (iINPC) and induced radial glial cells (iRGCs) (Figure 1C). Approximately 74% of the total cells were identified as radial glial cells, neuron-like cells and neuronal progenitors, while about 19% and 7% were oligodendroglial lineage cells (iODCs, iOPCs and iPPCs) and ciliated ependymal cells, respectively (Figure 1C). The sample identities were verified by checking genotype concordance between previously published whole-genome sequencing (WGS) data[32] and our generated scRNA-seq data (Figure 1D and Table S2). Subsequently, we compared the ICC-based estimation of MBP^+^ cells (range: 15%-40%) with the percentage of positive cells obtained from scRNA-seq data (range: 4%-32% and range: 16%-32% without sample 8) and found a similar distribution (Figure 1E), with a significant overall correlation coefficient (Spearman’s R = 0.69, Table S1). Of note, Sample 8 had the lowest percentage of MBP^+^ cells (4%) and the fewest reads (Table S1) as well as showed no contribution to the iODCs cell type (Figure 1F and Table S3), and was therefore excluded from the downstream analysis. Clusters were annotated based on the expression of known cell type markers: iODC (marked by *CNP, PLP1, MOG, MAG, MBP*), iOPC (marked by *PDGFRA*), iPPC (marked by *PDGFRA, MKI67 and TOP2A*), iCEP (marked by *SHH and CLDN5*), iNL2 (marked by *SNAP25, SNCA, TH and ZCCHC12*), iNL1 (marked by *SNAP25, SLC17A6 and RBFOX3)*, iINPC (marked by *HES6 and BTG2*) and iRGC (marked by *NES and GFAP*) (Figure 1G and Table S4). Notably, *LRRK2* demonstrated elevated expression levels in the iOPCs, iPPCs as well as in the iCEPs, consistent with earlier findings by Wang and colleagues[37], who observed high *LRRK2* expression in OPCs, microglia (cell type not present in our data) and endothelial cells from postmortem human substantia nigra single-nuclei RNA-seq data. Conversely, as reported by Wang and colleagues[37], *RIT2*, a PD susceptibility gene, was expressed not only in neuronal clusters but also in iOPCs. Integration of scRNA-seq data using Multi-marker Analysis of GenoMic Annotation (MAGMA_Celltyping) with PD genome-wide association studies (GWAS) showed a significant association (FDR adjusted P < 0.05) of iODCs, iOPCs and iNL2 with PD-linked risk loci (Figure 1H and Table S5). These results provide a cellular context that is appropriate for modeling the genetic risk in PD.

The *PDGFRA* and *LRRK2* genes were expressed in both iOPCs and iPPCs (Figure 2A). However, while *LRRK2* expression was also widespread in progenitors and iCEPs, *PDGFRA* expression was more specifically restricted to iOPCs and iPPCs (Figure 2A). The sub-clustering of iPPCs revealed three subtypes: iPPC_0, iPPC_1, and iPPC_2 (Figure 2B). As expected, all three subtypes showed significant expression of a proliferating marker *MKI67*. *PDGFRA* was expressed only in iPPC_0 and iPPC_2, indicating that these are OPC-like cells in a cycling state (Figure 2C and Table S4). Next, we assessed the similarity between the marker gene signatures of our cultured oligodendroglial lineage cells and those from postmortem human substantia nigra single-nuclei RNA-seq data by Wang and colleagues[37] (Figure 2D). Indeed, as anticipated, the iODCs clustered with human tissue ODCs, while the iOPCs clustered with human tissue OPCs (Figure 2D). This high concurrence between our iPSC-derived oligodendroglial cells and post-mortem cells further strengthens the use of these cells as models. Apart from iODCs, iOPCs, and iNL2, which were previously associated with risk loci, MAGMA analysis following sub-clustering identified a significant association of iPPC_2 with PD GWAS (Figure 2E and Table S5). Then, we analyzed differences in cell type proportions using a robust method, *speckle*[38], that accommodates variation in biological replicates (see Methods). We observed that all three cell types that exhibited significant *LRRK2* expression were enriched in PD lines compared to the controls, with p-values of 0.03 for iPPC_2, 0.05 for iCEP, and 0.07 for iOPC (Figure 2F and Table S6).

**Figure 2:**
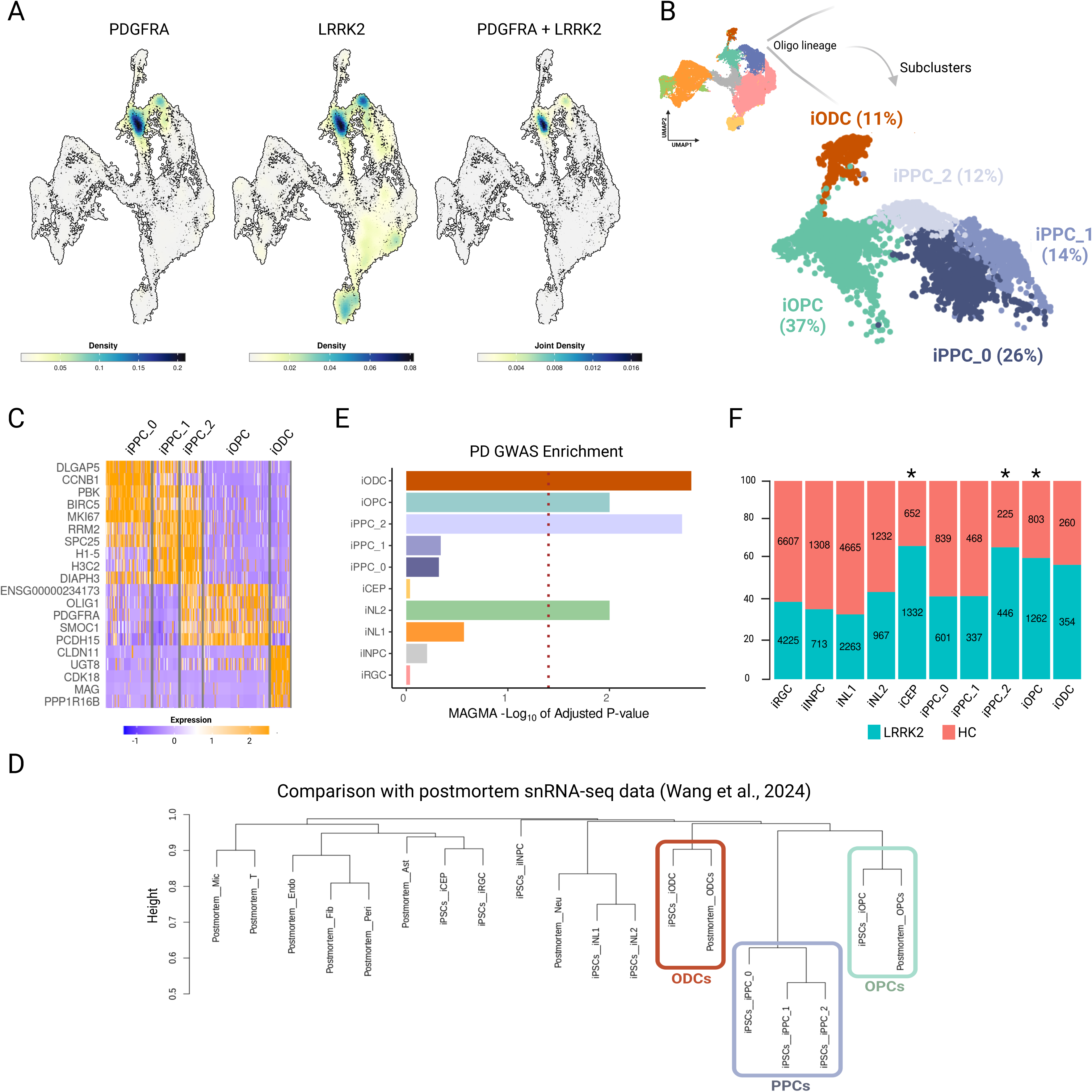
Sub-clustering of induced proliferative progenitor cells (iPPC). **(A)** Density plot representation of *PDGFRA* and *LRRK2* expression on UMAP visualization of cell types. **(B)** UMAP visualisation of cells of oligodendroglial lineage with sub-clustering analysis of induced-proliferative progenitor cells (iPPC). Percentages were calculated based on the total number of oligodendroglial lineage cells (5,595 cells). **(C)** Heatmap of the top markers for the iPPC clusters, iOPCs and iODCs **(D)** Clustering of cell type identities in comparison with postmortem snRNA-seq data from Wang et al.[37] **(E)** MAGMA gene set enrichment with iPPC clusters showed significant associations with induced-oligodendrocytes (iODCs), induced-oligodendrocyte precursor cells (iOPCs), induced-Neuron-like 2 (iNL2) cells and induced-proliferative progenitor cell 2 (iPPC_2) **(F)** Distribution of cell-type proportion with speckle analysis for statistical testing for differences in cell type composition; (* represents p-value < 0.1).

Differential gene expression analysis identified a total of 1,380 genes (795 unique genes) up-regulated in PD cell lines (Figure 3 and Table S7), whereas 3,770 genes (2,870 unique genes) were down-regulated (Figure 4 and Table S7). The significantly higher number of down-regulated genes compared to up-regulated ones was attributed to the iCEP and iNL1 clusters. Consequently, most up-regulated genes were uniquely present in one cell type (Figure 3A), while most down-regulated genes overlapped with iCEP or iNL1 (Figure 4A). Nevertheless, we observed a higher number of down-regulated genes compared to up-regulated genes in iODCs, iOPCs, and iPPC_0 (Figure 3A and Figure 4A). Conversely, in the iPPC_1 and iPPC_2 clusters, there were fewer down-regulated genes compared to up-regulated genes. Gene Ontology (GO) analysis revealed that up-regulated genes in iODCs and iOPCs were associated with negative chemotaxis and axon guidance related terms (Figure 3B and Table S8). Intriguingly, both iOPCs and iPPC_2 clusters displayed “Regulation Of Calcium-Mediated Signaling” biological process term due to the up-regulation of *RIT2*, *LRRK2* and *PLCG2*. Following this clue, we found that *LRRK2* was significantly up-regulated in iOPCs, iPPC_2 as well as iPPC_1 and iPPC_0 (Figure 3C). Notably, the Alpha-Synuclein gene (*SNCA)* was specifically up-regulated in the iNL2 cluster, which was characterized by the expression of dopaminergic neuron markers such as *TH* and *ZCCHC12* (Figure 3D and Figure 1G) and displayed cholesterol process related terms. On the other hand, genes associated with the “Semaphorin-Plexin Signaling Pathway” were down-regulated in iNL2 (*NELL1* and *NELL2*) and iOPCs (*NELL2, SEMA6A* and *PLXNA2*) (Figure 4B and Table S8). *SEMA6A* and *PLXNA2* were identified as specifically down-regulated in iOPCs (Figure 4C). Additionally, terms related to fatty acid and lipid transport were enriched in iOPCs and iPPC_2 down-regulated genes, while iPPC_1 showed enrichment of “Positive Regulation of T Cell-Mediated Immunity”. The down-regulated genes in iPPC_0 cluster exhibited the distinct terms related to cilia movement. Finally, we found down-regulation of myelin assembly-related genes in iODCs (Figure 4B and Table S7-8). For instance, *TPPP*, a modulator of microtubule dynamics and essential for myelination, and *MOBP*, a key regulator of myelin assembly, were specifically down-regulated in iODCs (Figure 4D).

**Figure 3:**
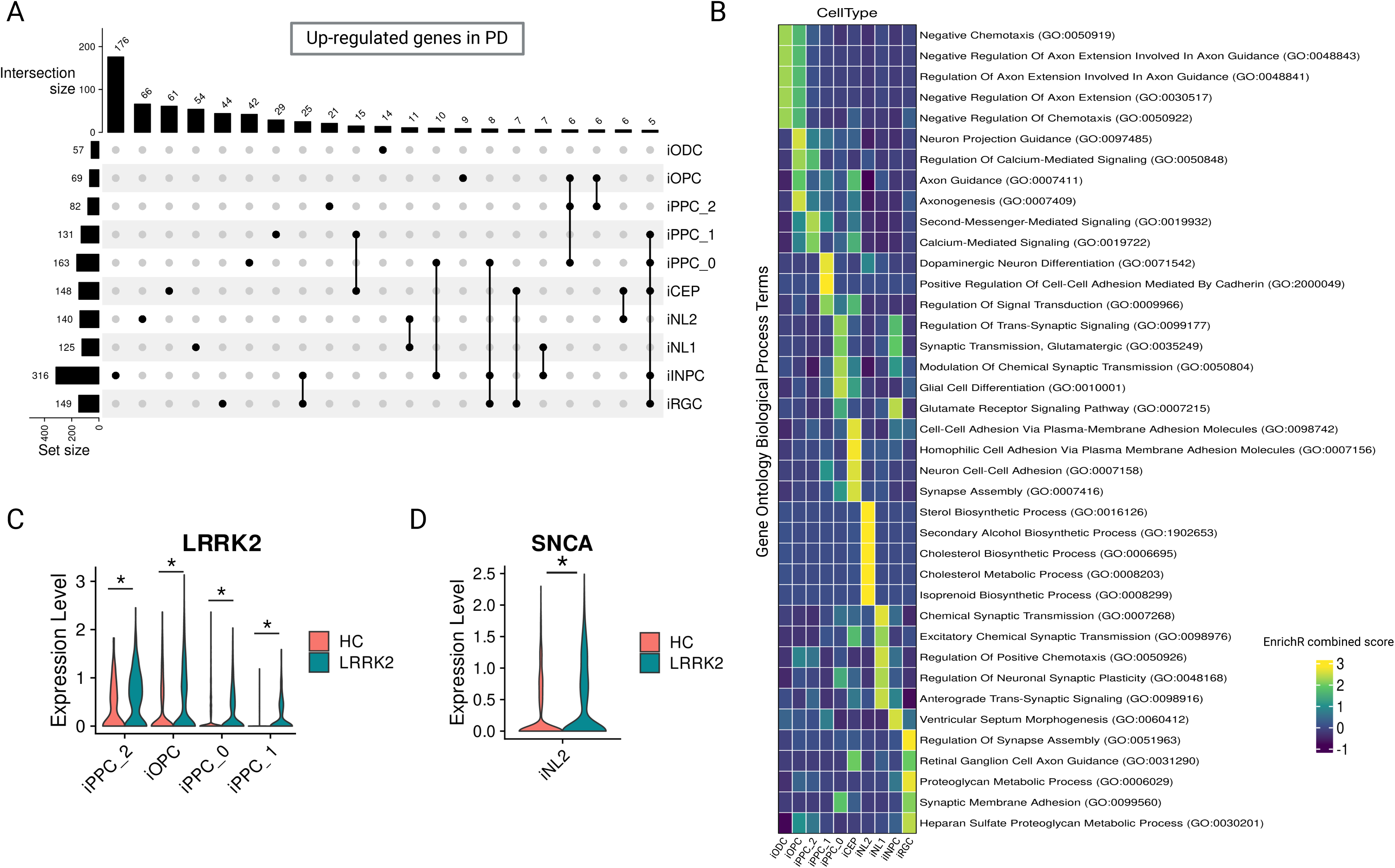
Differentially expressed up-regulated genes in each cell-type. **(A)** UpSet plot representing up-regulated differentially expressed genes in each cell type and the number of intersections between cell types with equal or more than 5 DEGs in common. **(B)** Gene ontology enrichment analysis of up-regulated genes. Top five biological process terms for each gene list indicated. Enrichr combined score is calculated by the logarithmic transformation of the p-value obtained from Fisher’s exact test, multiplied by the z-score representing the deviation from the expected rank. **(C)** Violin plot representing the gene expression level of *LRRK2* in HC and PD lines for the oligodendroglial lineage clusters. **(D)** Violin plots representing the gene expression level of *SNCA* in HC and PD lines for the iNL2 cluster (* represents adjusted p-value < 0.05).

**Figure 4:**
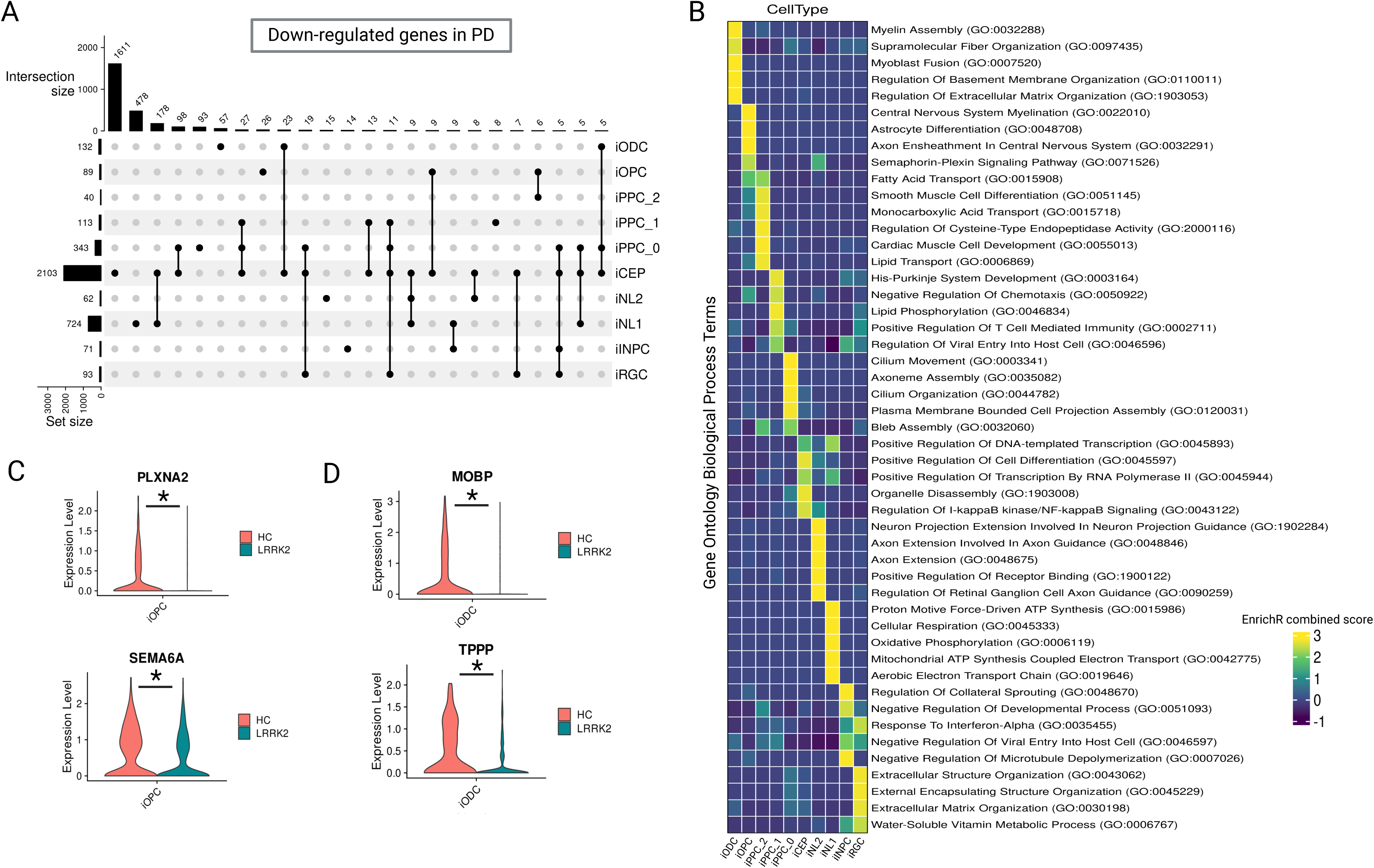
Differentially expressed down-regulated genes in each cell-type. **(A)** UpSet plot representing down-regulated differentially expressed genes in each cell type and the number of intersections between cell types with equal or more than 5 DEGs in common. **(B)** Gene ontology enrichment analysis of down-regulated genes. Top five biological process terms for each gene list indicated. Enrichr combined score is calculated by the logarithmic transformation of the p-value obtained from Fisher’s exact test, multiplied by the z-score representing the deviation from the expected rank. **(C)** Violin plots representing the gene expression level of *PLXNA2* and *SEMA6A* in HC and PD lines for the iOPC cluster. **(D)** Violin plots representing the gene expression level of *MOBP* and *TPPP* in HC and PD lines for the iODC cluster (* represents adjusted p-value < 0.05).

Given the observed dysregulation in several signaling pathways, we hypothesized that this could result in significant signaling changes between cells across PD and HC lines. To investigate this, we used CellChat (v2)[39] to predict the main signaling inputs and outputs for cells under each condition and subsequently identified key signaling alterations between the PD and HC lines. We compared the information flow of each signaling pathway between PD and HC lines, defined as the sum of communication probabilities among all cell group pairs in the inferred network (Figure 5A and Table S9). Cell-cell communication analysis revealed significant changes in several signaling pathways, including the deactivation of PSAP signaling (Figure 5B) and the activation of MIF signaling (Figure 5C) in PD lines. Notably, PSAP signaling was concentrated in iODCs, and *PSAP*, a lysosome-related gene, has been recently associated with PD risk[40]. On the other hand, concerning macrophage migration inhibitor factor (MIF) signaling (turned on in PD lines here), Park and colleagues[41] recently demonstrated that the pathologic α-synuclein neurodegeneration in PD is mediated through MIF nuclease activity, and genetic depletion of MIF in mice prevents the loss of dopaminergic neurons. Additionally, we found the largest difference in information flow within the SEMA6 signaling network, which was overall increased in PD lines (Figure 5D and Table S9). However, iOPCs derived from these PD lines exhibited a significant loss of information flow within the SEMA6 signaling pathway (Figure 5D, right panel).

**Figure 5:**
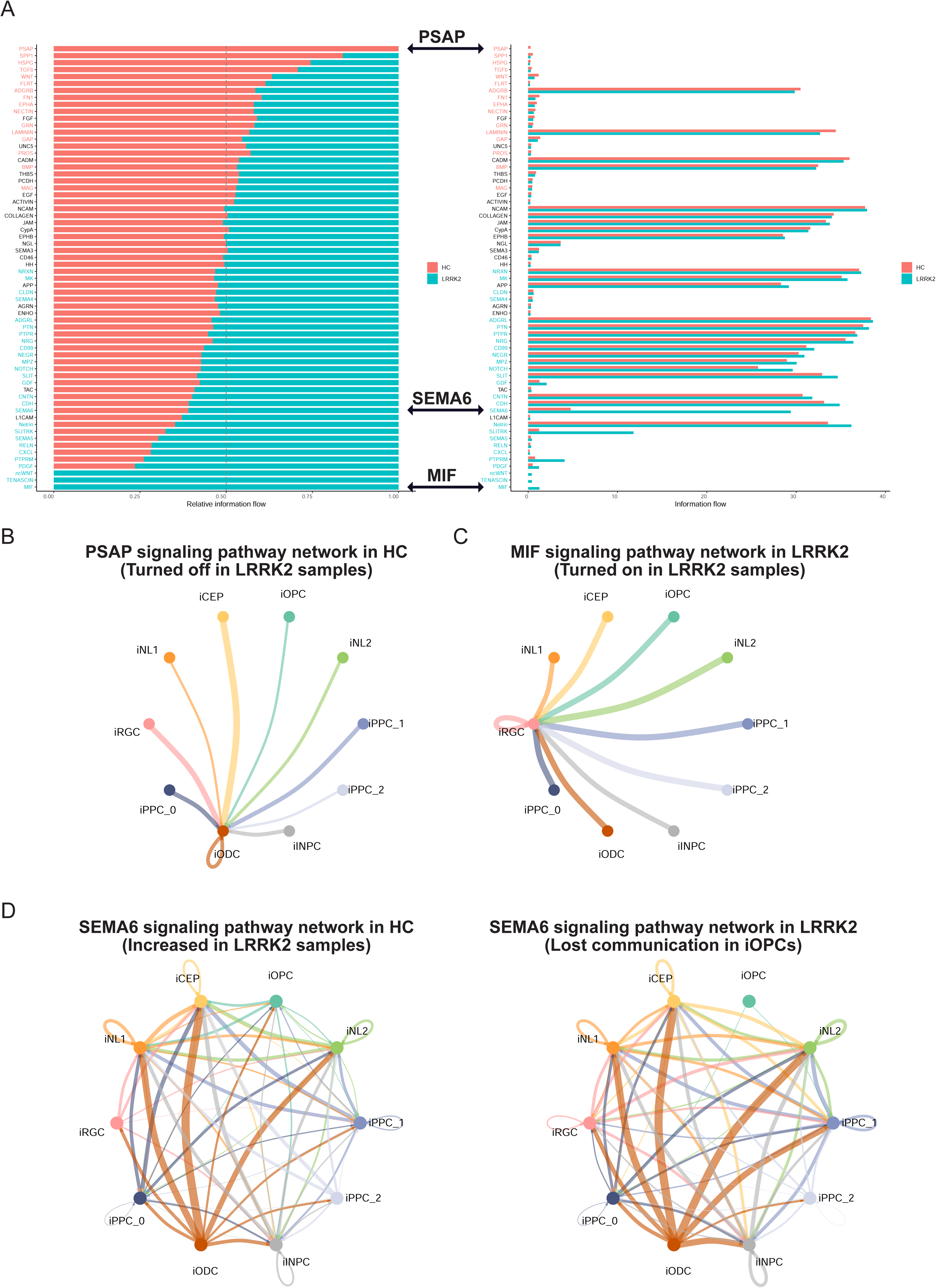
Cell-cell communication analysis. **(A)** Table representing signals that contribute to outgoing and incoming signals for HC and LRRK2 PD lines. **(B)** The chord circle plot displays significantly interacting pathways and communication probabilities of PSAP signaling, which is present only in HC **(C)**, MIF signaling, which is present only in PD lines **(D),** and Left: SEMA6 signaling in HC Right: SEMA6 signaling in PD lines.

An unsupervised pseudotime analysis was conducted on oligodendroglial lineage cells, predicting a trajectory that spans from cycling cells of iPPCs to mature iODCs (Figure 6A). A total of 183 genes were identified that were significantly associated with pseudotime progression (FDR adjusted P < 0.05) and also differentially expressed between conditions (FDR adjusted P < 0.05) (Figure 6B and Table S10). Gene Ontology analysis revealed that these genes were enriched for terms related to semaphorin-plexin signaling, negative chemotaxis, cell migration and development (Figure 6C and Table S10). We noted increased expression of *SHH* at earlier pseudotime stages, along with higher expression levels in PD lines (Figure 6B), aligns with findings by Schmidt and colleagues[42], who reported that shortened primary cilia are associated with increased Sonic Hedgehog (SHH) signal transduction. We summarized the expression changes of genes related to the hedgehog and semaphorin-plexin signaling pathways in oligodendroglial lineage cells (Figure 6D). *SHH* and *SEMA6D* were up-regulated in iPPC_1 and iPPC_0, whereas *GLI* genes and *SEMA3A* were down-regulated. *SEMA6A* and *PLXNA2* were specifically down-regulated in iOPCs. To validate our findings and extend our understanding of *SEMA6A* and *PLXNA2* expression in OPCs, we reanalyzed publicly available single-nuclei RNA sequencing data from human postmortem prefrontal cortex and anterior cingulate brain regions (Dehestani 2023). As presented in Figure 6E, *SEMA6A* and *PLXNA2* displayed a down-regulation in OPCs of LRRK2 brain samples compared to controls. Although both genes showed a down-regulation pattern, it is important to note that only *SEMA6A* reached statistical significance (log_2_ FC = -0.32, adjusted P = 1.46e-06).

**Figure 6:**
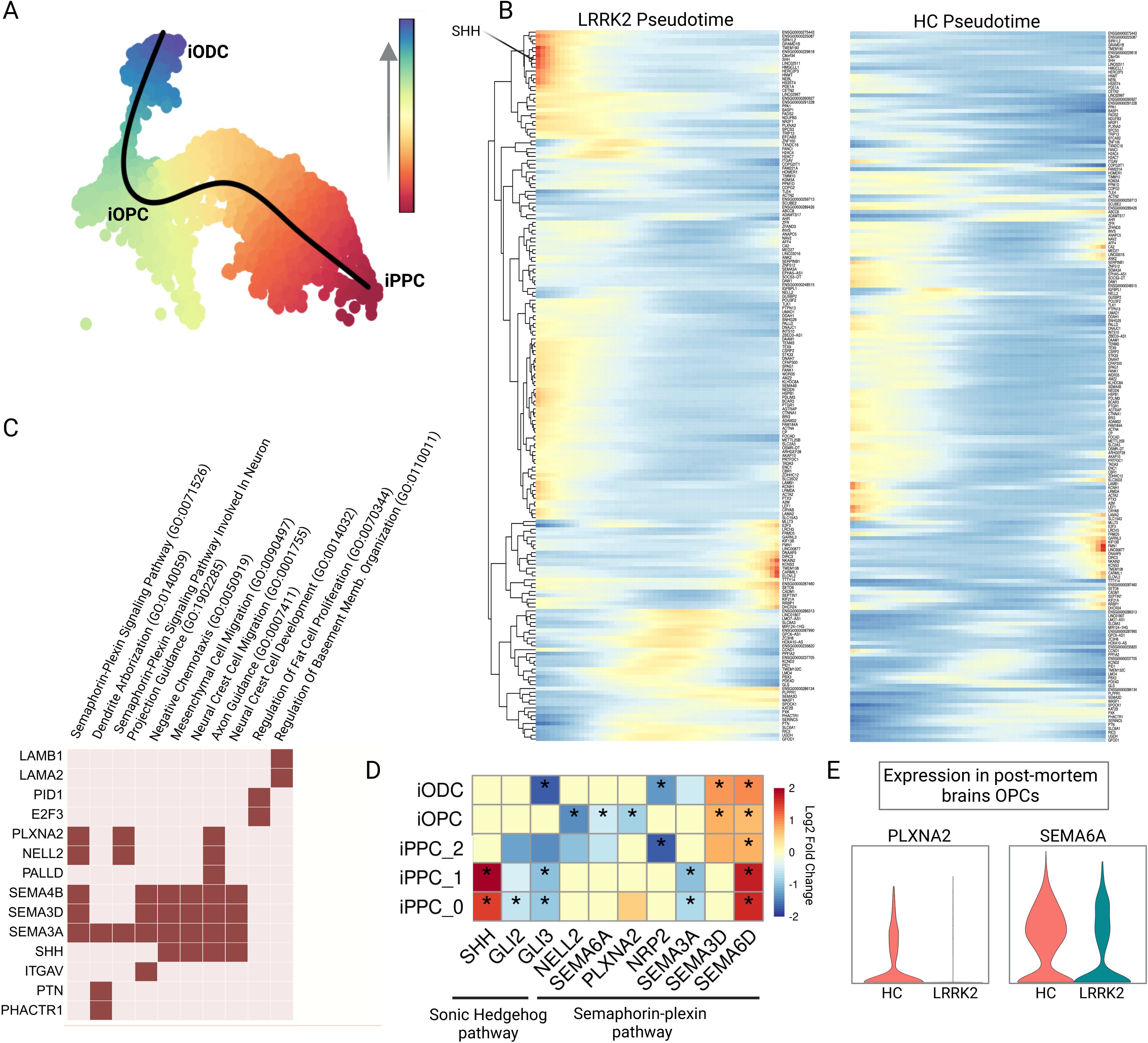
Single-cell trajectory and pseudotime analysis. **(A)** Pseudotime analysis for the oligodendroglial lineage (cell types iODC, iOPC, and iPPC). **(B)** Heatmap of genes overlapping between significant pseudotime-associated and condition test gene lists. **(C)** Gene ontology enrichment analysis of pseudotime-associated genes using biological process terms. **(D)** Heatmap showing differential expression of genes associated with semaphorin-plexin and sonic hedgehog pathways (* represents FDR-corrected p-value < 0.05). The semaphorin-plexin and sonic hedgehog related genes were extracted from CellChat signaling pathways using plotGeneExpression function. Additionally, GLI genes (*GLI1, GLI2, GLI3*) were included. Only the genes that were significantly differentially expressed in at least one cell type are displayed. **(E)** Violin plots showing the expression of *SEMA6A* and *PLXNA2* in postmortem brain OPCs with and without LRRK2-G2019S mutation. The snRNA-seq data was reanalysed from previous publication[13] (GSE272760) using a bioinformatics pipeline from this study.

Mutations in LRRK2 have previously been shown to cause primary cilia defects in striatal cholinergic interneurons and astrocytes[43]. In line with this, we observed down-regulation of cilia genes in iPPC clusters (Figure 4B). Since *LRRK2* expression levels were found to be highly expressed in iOPCs (Figure 1G), we investigated whether the same LRRK2 mutation could play a role in OPC ciliation. Firstly, day 11 iOPCs were strongly positive for the ciliary marker Arl13b (Figure 7A), clearly visible in cells stained for platelet derived growth factor alpha (PDGFRα), a marker used to identify OPCs in the brain[44,45]. This is in agreement with the previous report on the presence of primary cilia in OPCs[46]. In contrast, the percentage of cilia-positive cells was lower in iOPCs harboring the LRRK2 p.G2019S mutation (Figure 7B and C). Shortened cilia have previously been associated with higher *SHH* expression[42,47], which was also observed in our pseudotime analyses of oligodendroglial lineage in LRRK2 p.G2019S PD lines during early stages of oligodendroglial differentiation (Figure 6B), as cilia is absent in mature oligodendrocytes[46].

**Figure 7:**
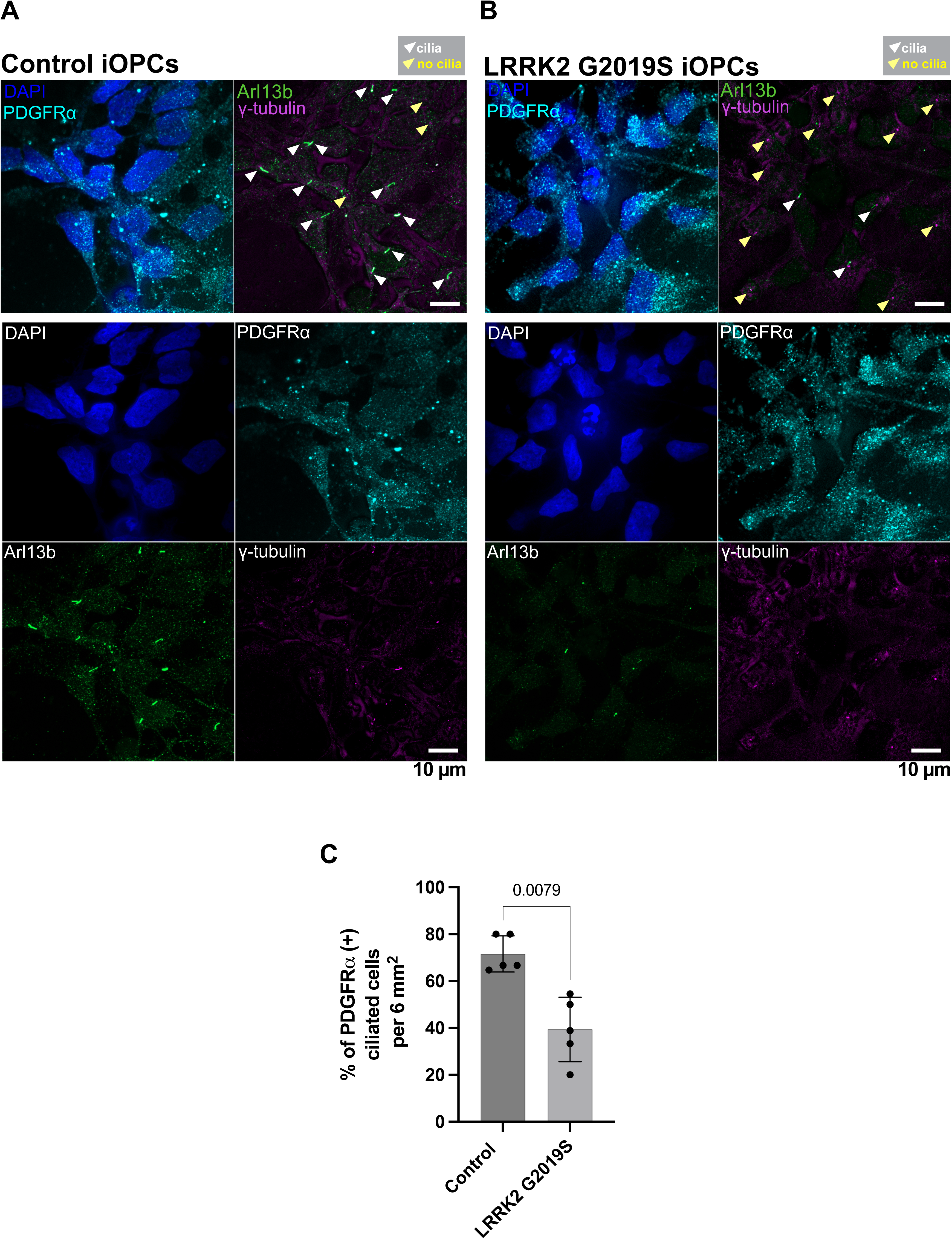
Reduced ciliation in LRRK2-G2019S iOPCs. **(A-B)** Airyscan fluorescence images of control (A) and LRRK2 p.G2019S (B) iOPCs (day 11) immunolabeled with antibodies directed against OPC marker PDGFRα (cyan), ciliary marker Arl13b (green) and centrosomal marker γ-tubulin (green). DAPI was used to label the nuclei. Single channel images are shown beneath each merged panel (Scale bars, 10 µm). **(C)** Percentage of PDGFRα-positive cells with cilia (mean ± S.D.) from 6 mm2 areas on coverslips containing n ≥ 5 cells per area.

## Discussion

In this study, we provide novel insights into the cellular and molecular mechanisms through which the LRRK2 p.G2019S mutation contributes to PD pathogenesis, with particular emphasis on oligodendroglial lineage cells. We used well-characterized PPMI human iPSC lines derived from PD patients with the LRRK2 p.G2019S mutation and from healthy controls, and generated scRNA-seq data from oligodendroglial lineage cells derived from these lines. We discovered significantly high expression of *LRRK2* in OPCs, proliferating OPCs as well as ependymal cell clusters compared to other cell types. All three cell types showed a significant increase in cell numbers in PD lines relative to controls, indicating a direct influence of LRRK2 on cell proportions. The up-regulation of *LRRK2* in PD OPCs and proliferating OPCs, along with *RIT2* (a PD risk gene[48]) and *PLCG2* (a phagocytosis- and immune-related gene[49]), consequently resulted in the enrichment of calcium-mediated signaling GO term. Previous studies have demonstrated that *LRRK2* overexpression increases autophagosome[50] numbers via activation of a calcium-dependent pathway, potentially influencing the localization of α-synuclein[51]. Since *LRRK2* is not expressed in neurons here as compared to OPCs, it is tempting to speculate that LRRK2 in OPCs may interact with SNCA in neurons by dysregulated calcium fluxes.

We identified dysregulated genes related to cilia movement, SHH signaling and semaphorin-plexin pathways in oligodendroglial lineage cells. Recent studies from various laboratories suggest that impairments in primary cilia and SHH signaling likely contribute to PD pathology[42,47,52–55]. Importantly, the convergence on impaired ciliogenesis and SHH signaling in PD, incorporates data from both genetic and sporadic forms of the disease and is consistent across multiple tissues and iPS-derivation protocols. It has been proposed that *LRRK2* mutations affect the ability of cells to respond to cilia-dependent SHH signaling. Consistent with these observations, we identified a subcluster of proliferating OPCs that exhibited down-regulation of cilia-related genes, while *SHH* expression was significantly up-regulated. Intriguingly, Pinskey and colleagues[56] demonstrated that the GTPase-activating protein domain of plexin is necessary to enhance Hedgehog signaling at the level of GLI transcription factors, and this enhancement depends on intact primary cilia. Semaphorin-plexin pathways might be the missing link between primary cilia and SHH signaling. We observed a complex dysregulation of the genes in the semaphorin-plexin pathway: there was an overall increase in SEMA6 signaling communication in PD lines, yet OPCs derived from these lines specifically lost SEMA6 signaling. It is known that the migration of OPCs is modulated by a balance of effects mediated by members of the semaphorin and netrin families[57] and Sema3A is known to be a potent, selective, and reversible inhibitor of OPC differentiation *in vitro*[58]. We found that *SEMA3A* and cilia-related genes are down-regulated in the iPPC_0 cluster, which may contribute to cilia impairments in later-stage iOPCs. A recent study[46] in the mouse primary visual cortex found that OPCs possess primary cilia. Whether human OPCs require intact primary cilia for semaphorin-plexin and SHH signaling is yet to be resolved. Another explanation for the overall increase in semaphorin-plexin-related genes and *SHH* is their involvement in cell migration[59,60], potentially as a response to the down-regulation of myelin assembly-related genes in ODCs, with cilia impairments possibly hindering this migration. Alternatively, the specific down-regulation of *SEMA6A* in PD OPCs might delay oligodendrocyte differentiation, leading to a reduced expression of myelin assembly related genes in ODCs[61] from PD lines.

Our study has some limitations due to the shortcomings of iPSC models and the availability of limited resources. In general, iPSC lines and differentiation protocols exhibit variability, and the transcriptional profiles of iPSC-derived oligodendroglial cells may not mirror those of mature or aged cell types. Additionally, RNA expression levels might be underestimated due to dropout effects in scRNA-seq and may not accurately reflect protein abundance. Moreover, cell composition analysis using droplet-based scRNA-seq data can be sensitive to cell size, cell shape, and subsequent computational pipelines. Therefore, we strongly recommend interpreting our scRNA-seq cell composition analysis results with caution. For instance, ICC indicated that approximately 55% of cells were O4 positive (including oligodendrocytes with myelin sheaths), whereas scRNA-seq showed only 19% oligodendroglial lineage cells, emphasizing the importance of using ICC in conjunction with scRNA-seq. This discrepancy suggests the need for single-nuclei and/or full-length RNA sequencing. We compared the iPSC-derived cell types generated in this study with those identified in postmortem brain samples and found transcriptional agreement in their identities. However, we cannot rule out the possibility of missing rare subpopulations of OPCs and ODCs cell types[62–64] present in human brains that are susceptible to aging and aging-associated diseases like PD. Furthermore, we recognize that a larger cohort of iPSC lines may be necessary to detect subtle molecular changes in disease mechanisms. Nevertheless, iPSC technology has evolved into a crucial asset for investigating the mechanisms involved in PD pathogenesis and for conducting preclinical exploration aimed at evaluating potential treatments[65].

Altogether, we present the first scRNA-seq and immunocytochemistry data from oligodendroglial lineage cells derived from PD patient-iPSCs with LRRK2 p.G2019S mutation. We propose that dysfunctional semaphorin-plexin signaling, along with cilia movement and SHH signaling, might represent early events in PD pathology. The findings highlight the need for a deep exploration of the complex interactions among semaphorin-plexin, sonic hedgehog and cilium pathways in PD. Future studies involving other PD-relevant mutations, along with isogenic controls, will help elucidate the common and distinct pathways of PD pathoetiology in oligodendroglial cells. We envision that our work will serve as a valuable resource for uncovering potential targets in PD.

## Methods

### Culturing of induced pluripotent stem cells (iPSCs)

The Parkinson’s Progression Markers Initiative (PPMI) iPSCs included in this study have previously been generated and characterized[32]. Human iPSCs were cultured as colonies in feeder-free conditions using Essential 8 Flex Medium (Thermo Fisher Scientific, A2858501) on Matrigel hES-qualified (Corning, 354277) coated plates. Colonies were passaged as aggregates every 5-6 days. Briefly, iPSC colonies were treated with Gentle Dissociation Reagent (GDR, StemCell technologies, 07174) for 7 min at room temperature. GDR was then replaced with E-8 flex media and colonies were gently triturated by pipetting three times to obtain a suspension of small aggregates. Small colonies were plated onto Matrigel-coated plates, and the media was refreshed every other day.

### Differentiation of iPSCs into neural progenitor cells (NPCs)

The small molecules NPCs (smNPCs) were derived from iPS cells using the method described in Reinhardt et al. 2013[35], applying a few adjustments as described in Dhingra et al, 2020[66]. The smNPCs were cultured on Matrigel (Corning, 354234) coated plates in the expansion medium (N2B27) supplemented with 3 µM CHIR (R&D System, 4423), 0.5 µM PMA (Cayman Chemical, 10009634) and 64mg/l Ascorbic acid (AA, Sigma-Aldrich, A8960) for up to 8 passages before use in any downstream experiments. N2B27 medium consisted of DMEM/F-12-GlutaMAX (Thermo Fisher Scientific, 31331093) and Neurobasal (Thermo Fisher Scientific, 21103049) mixed at 1:1 ratio, supplemented with 1:200 N-2 supplements (Thermo Fisher Scientific, 17502048), 1:100 B-27 supplements without vitamin A (Thermo Fisher Scientific, 12587010), 1:200 GlutaMAX™ (Thermo Fisher Scientific, 35050), 5 μg/ml insulin (Sigma-Aldrich, I9278), 1:200 non-essential amino acids (NEAA, Thermo Fisher Scientific, 11140050), 55 μM 2-mercaptoethanol (Thermo Fisher Scientific, 21985023), and 1:100 penicillin/streptomycin. The smNPCs were routinely passaged using Accutase (Sigma-Aldrich, A6964).

### Differentiation of smNPCs into oligodendroglial-lineage cells

For the differentiation of small molecule neuronal precursor cell (smNPCs) into viral induced oligodendrocytes (hiOL), 80-90% confluent cells were detached with accutase (Pan Biotech, #P10-21100) and seeded in smNPC medium in CultrexTM (R&D systems #3532-005-02) -coated 6-well plates with a density of 3*105 cells per well. The next day, cells were transduced with the SON-Puro lentivirus and were incubated overnight before washing on the next day. A day after washing, the differentiation was started by changing the medium to glial induction medium (GIM). For selection 2 µg/ml puromycin (Gibco, #A11138-03) was added from differentiation day 2 to day 7. Medium was changed to glial differentiation medium on day 4 of the differentiation and was changed every other day. To obtain a pure culture cells were FACS sorted on day 21 of differentiation for O4-APC (#130-118-978, Miltenyi Biotech). After the FACS sort cells were cultivated in GDM for another 14 days and medium was changed every other day. At day 35 of the differentiation cells were fixated with 4%PFA.

### Immunocytochemistry (ICC)

For ICC, cells were cultured on CultrexTM (R&D systems #3532-005-02) -coated 48-well or 96-well plates. After fixation with 4% PFA for 20 min at RT, cells were washed twice with PBS and either stored at 4 °C or used immediately for staining. For blocking of non-specific binding sites, cells were incubated 45 min in blocking buffer, containing PBS with 5% normal goat serum (NGS) and 5% fetal bovine serum (FBS), at RT. For intracellular binding epitopes, cells were subsequently permeabilized with 0.5% Triton-X-100 in PBS for 5-7 min at RT. Afterwards, primary antibodies (Table S11) were diluted in the blocking buffer and were applied. Cells were incubated at 4 °C overnight. The next day, cells were washed twice with PBS, Cy-3 or Alexa-488 conjugated antibodies (Table S11) were applied and incubated for 1 h at RT in the dark. For double stainings steps for applying primary and secondary antibodies were repeated. After washing cells twice with PBS, to counterstain the nuclei 1µg/mL 4’-,6-diamidino-2-phenylindole (DAPI) in PBS was added for 10 min followed by additional three washing steps. To visualize fluorescent staining cells were analyzed by an Olympus CKX53 inverted microscope (Olympus). For each cell line, 10 images of a technical triplicate were taken and the mean value determined.

### Fluorescence associated cell sorting (FACS)

To prepare cells for the FACS procedure, cells were detached with accutase (Pan Biotech, #P10-21100) 10 min at 37 °C. Afterwards cells were resuspended and diluted in a ratio of 1:10 in split medium and centrifuged for 5 min at 200 rcf. After centrifugation, cells were resuspended in FACS buffer, which consists of PBS with 0.5% BSA Fraction V (Gibco, #15260037). Cell numbers were determined using a Neubauer counting chamber. Cells were stained with the O4-APC antibody (Miltenyi Biotech, #130-118-978) according to the manufacturer’s manuscript. For FACS sorting, cells were stored in ice.

### Single-cell transcriptome library preparation and sequencing

Single-cell suspension concentration was determined by automatic cell counting (DeNovix CellDrop, DE, USA) using an AO/PI viability assay (DeNovix, DE, USA) and counting cells as dead cells. Single-cell gene expression libraries were generated using the 10x Chromium Next gel beads-in-emulsion (GEM) Single Cell 3’ Reagent Kit v3.1 (10x Genomics, CA, USA) according to manufacturer’s instructions. In brief, cells were loaded on the Chromium Next GEM Chip G, which was subsequently run on the Chromium Controller iX (10x Genomics, CA, USA) to partition cells into GEMs. Cell lysis and reverse transcription of poly-adenylated mRNA occurred within the GEMs and resulted in cDNA with GEM-specific barcodes and transcript-specific unique molecular identifiers (UMIs). After breaking the emulsion, cDNA was amplified by PCR, enzymatically fragmented, end-repaired, extended with 3′ A-overhangs, and ligated to adapters. cDNA QC and quantification was done with Tape Station High Sensitivity D5000 ScreenTape (Agilent, CA, USA). P5 and P7 sequences, as well as sample indices (Chromium Library Construction Kit, 10x Genomics, CA, USA) with Dual Index Kit TT Set A, (10x Genomics, CA, USA), were added during the final PCR amplification step. The fragment size of the final libraries was determined using the Bioanalyzer High-Sensitivity DNA Kit (Agilent, CA, USA). Library concentration of the final libraries was determined using the Tape Station High Sensitivity D1000 ScreenTape (Agilent, CA, USA). Single-cell RNA libraries were pooled and paired-end-sequenced on the Illumina NovaSeq 6000 platform (Illumina, CA, USA), using a S1 flow cell.

### Single-cell RNA-seq quality control and analysis

The sequencing reads were demultiplexed using the 10x Genomics Cell Ranger v8.0 and to perform alignment against the 10x Genomics pre-built Cell Ranger reference GRCh38-2024-A to generate count matrices for each sample. Seurat[67] objects v5.0.3 were created for each sample with a cut-off value of 200 unique molecular identifiers (UMIs) expressed in at least 3 cells. Cells with fewer than 1000 genes detected or more than 5% of reads mapping to mitochondrial genes were removed. Each sample was processed individually, and doublets and multiplets were filtered out using DoubletFinder v2.0.3[68] with recommended settings. Following the filtering steps applied, the dataset contained 35184 high-quality cells that were used for the following analyses. The individual Seurat objects were merged and normalized by the SCTransform[69] method in Seurat v5.0.3 with mitochondrial reads regressed out. Principal component analysis (PCA) was then performed on the normalized data using the ‘RunPCA’ function. Integration and batch correction across samples were achieved using the Harmony R package version 1.2.0 with the ‘HarmonyIntegration’ function. The first 30 principal components (PCs) from the Harmony-corrected data were used as inputs for downstream analyses. A shared-nearest-neighbor graph was constructed using the ‘FindNeighbors’ function and cells were clustered using the Louvain algorithm implemented in the ‘FindClusters’ function with a resolution of 0.1. Cluster 2 was then further sub-clustered using ‘FindSubCluster’ in order to separate the iOPC cluster and the iINPC cluster using the Louvain algorithm with a resolution of 0.1. The cell identities were saved in the metadata in the column ‘BroadCellType’ (Table S3). Sub-clustering was performed on the iPPC cluster using the Louvain algorithm with a resolution of 0.15 using the ‘FindSubCluster’ function. The sub-clustered cells were assigned new identities representing different iPPC cell types, and the metadata was updated accordingly under the column ‘CellType’. UMAP was performed to visualize the clusters in two dimensions using the ‘RunUMAP’ function. The Seurat function ‘FindAllMarkers’ with default parameters was used in order to identify markers for the clusters. Known markers based on previously reported literature were confirmed in order to determine the identity of each cluster (Table S4). The proportion of cell types was analyzed using the ‘speckle[38]’ R package v0.99.7 and the ‘propeller’ function with the ‘CellType’, ‘SampleID’ and ‘Mutation’ columns from the Seurat metadata as input (Table S6). To compare our data with postmortem brain tissues, ClusterMap[70] v0.1.0 was executed on the top 500 genes (sorted by p-value) from each cluster with at least one log fold-change, resulting in a total of 5,000 marker genes from this study and 3,289 marker genes from Wang et al[37].

### Genotype and sample identity check

We evaluated the correlation between single-cell sequencing data and donor (blood-derived) whole-genome sequencing (WGS) data to validate sample origin and identity in our scRNA-seq data. The subset-bam v1.1.0 was used to subset BAM files by extracting the alignment records with valid filtered cell barcodes generated by Cell Ranger (https://github.com/10XGenomics/subset-bam). The cellsnp-lite[71] v1.2.3 tool in Mode 2b was used in pseudo-bulk manner to call genotypes in scRNA-seq BAMs using the WGS VCF file from chromosome 12. The resulting VCF files from scRNA-seq data were indexed using bcftools[72] v1.20. Following the indexing, sample concordance between the WGS VCF and single-cell VCF files was checked using the ‘gtcheck’ function in bcftools. Finally, a summary table was created with the discordant rates. The discordance rate indicates the level of inconsistency between the samples, where a lower discordance rate signifies higher consistency between the WGS and single-cell data (Table S2).

### Differential expression and gene ontology enrichment analysis

Differential gene expression analysis was performed using the non-parametric Wilcoxon rank sum test implemented in the ‘FindMarkers’ function of Seurat v5.0.3 with default parameters with the exception of the parameters ‘min.pct’ and ‘min.diff.pct’ where a threshold of 0.1 was used. Differentially expressed genes between PD and HC were determined for each cell type. Gene ontology enrichment analysis for biological processes was performed using EnrichR[73] library GO_Biological_Processes_2023. Enrichr combined score is calculated by the logarithmic transformation of the p-value obtained from Fisher’s exact test, multiplied by the z-score representing the deviation from the expected rank (Table S7; DEGs, Table S8; GO).

### Cell-type association with genetic risk of PD

Association analysis of cell type-specific expressed genes with genetic risk of PD was performed as described previously[32], using Multi-marker Analysis of GenoMic Annotation (MAGMA) v2.0.2, in order to identify disease-relevant cell types in the data[74,75]. MAGMA, as a gene set enrichment analysis method, tests the joint association of all SNPs in a gene with the phenotype, while accounting for LD structure between SNPs. Competitive gene set analysis was performed on SNP p-values from the latest PD GWAS summary statistics including 23andMe data and the publicly available European subset of 1000 Genomes Phase 3 was used as a reference panel to estimate LD between SNPs. SNPs were mapped to genes using NCBI GRCh37 build (annotation release 105). Gene boundaries were defined as the transcribed region of each gene. An extended window of 10 kb upstream and 1.5 kb downstream of each gene was added to the gene boundaries.

### Cell-cell communication analysis

We used CellChat[39] v2.1.2 to perform a comparison analysis between LRRK2 and HC lines. As indicated by the vignette, we ran CellChat on each group (LRRK2 or HC) independently, and then merged the two CellChat objects. We used the functions computeCommunProb, filterCommunication (50 cells), computeCommunProbPathway, aggregateNet, and netAnalysis_computeCentrality with default parameters, utilizing CellChatDB.human as the ligand-receptor interaction database. The CellChat objects from LRRK2 and HC were merged using mergeCellChat function. To create Figure 5A and Table S9, the signaling networks were ranked based on the information flow using the rankNet function. We utilized the netVisual_aggregate function to visually compare cell-cell communication using a Circle plot as shown in Figure 5B-C.

### Single-cell trajectory and pseudotime analysis

For the pseudotime analysis and trajectory inference, we utilized Slingshot[76] v2.10.0. Our input data comprised a gene expression matrix extracted from a subset of fully processed Seurat object from previous steps, only including oligodendroglial lineage cells: iPPC_0, iPPC_1, iPPC_2, iOPC, and iODC. Slingshot operates by constructing a minimum spanning tree (MST) and subsequently fitting principal curves to the high-dimensional gene expression data. This approach effectively captures the underlying nonlinear relationships and delineates the structure of cellular differentiation and transitional trajectories. Importantly, it enables the inference of continuous cell progression through various developmental stages without the need to determine root cells explicitly (Figure 6A). Following trajectory inference, we used TradeSeq[77] v1.16.0 to find pseudotime-associated genes which are differentially expressed between conditions. It fitted a generalized additive model (GAMs) to the expression data of each gene along the pseudotime. GAMs are flexible models that can capture complex, non-linear relationships between gene expression and pseudotime. AssociationTest implemented in this tool was used to firstly identify pseudotime-associated genes. Subsequently, its conditionTest function was executed to find genes that are differentially expressed between conditions. We proceeded to look at the overlaps (n=183) among significant genes based on multiple testing adjusted p-values (<0.05) derived from both associated and differentially expressed gene lists. This analysis underscored distinct expression patterns among pseudotime-associated genes in LRRK2 and HC lines (Figure 6B). Finally, we utilized EnrichR to investigate the functional enrichment of these overlapping genes within biological processes. The semaphorin-plexin and sonic hedgehog related genes were extracted from CellChat signaling pathways using plotGeneExpression function. Additionally, GLI genes (*GLI1, GLI2, GLI3*) were included.

### Immunofluorescence and fluorescence microscopy for cilia

For cilia stainings, one line from each group (LRRK2 and HC) were randomly selected to differentiate (day 11) on glass slides. Cells were seeded on glass coverslips and fixed with 4% (v/v) paraformaldehyde in 1x phosphate-buffered saline (PBS) for 20 min followed by three washes in PBS. Cells were permeabilized with 0.5% (v/v) Triton X-100 in PBS for 10 min followed by three washes in PBS. They were further blocked for 30 min in 5% bovine serum albumin (BSA, Sigma-Aldrich) in PBS and then incubated overnight at 4 °C with the primary antibodies listed in Table S11. Subsequently, cells were washed with PBS thrice the following day and incubated with Alexa Fluor-conjugated secondary antibodies (Thermo Fisher Scientific) for 1 h at room temperature, followed by three washes in PBS. Coverslips were mounted on glass slides with ROTI®Mount FluorCare DAPI (Roth Carl) at 4°C. For imaging, The LSM 980 inverted confocal laser scanning microscope with Airyscan 2 (Carl Zeiss Microscopy) accompanied with 63×/1.40 numerical aperture (NA) plan-apochromat differential interference contrast (DIC) M27-Oil immersion objective and 32-channel gallium arsenide phosphide (GaAsP)-photomultiplier tubes (PMT) area detector with 405nm, 488 nm, 561 nm and 633 laser lines was used in this study. Images were acquired and processed using ZEN black imaging software (Zeiss).

### Image and statistical analysis

Images were pseudocolor-coded, adjusted for brightness and contrast, using the open-source image processing software FIJI (ImageJ)[78]. For ciliary analyses, Mann–Whitney test was used when comparing two datasets. Differences were accepted as significant for P < 0.05. Prism version 10 (GraphPad Software) was used to plot, analyze, and represent the data.

## Availability of data and materials

The supporting results are provided in supplementary tables. Scripts are available at https://github.com/nnkarma12/scRNAseq_iOligo.

## Authors’ contributions

V.B. conceived the project. E.F., T.G., T.K. and V.B. provided the resources. M.D., N.K., S.T. and V.B. performed the computational analysis of scRNA-seq. T.K. conceived the oligodendroglia differentiation experiments. W.K. performed the oligodendrocyte differentiation experiments, staining and image analysis. W.S. carried out the genotype check analysis. P.V., J.F. and N.M.R. performed the cilia stainings in iOPCs. A.D. and S.R.N differentiated the iPSCs to NPCs. J.T., D.S. and N.F. performed cell isolation and 10x Genomics library preparation. M.D., N.K. and V.B. wrote the manuscript. All authors contributed to and reviewed the manuscript.

## Acknowledgments

Patient-derived iPSC lines, reprogrammed from the peripheral blood mononuclear cells, were obtained from the Golub Capital iPSC Parkinson’s Progression Markers Initiative (PPMI) substudy (https://www.ppmi-info.org/access-data-specimens/request-cell-lines) and were also used in FOUNDIN-PD consortium (https://www.foundinpd.org/). PPMI – a public-private partnership - is funded by the Michael J. Fox Foundation for Parkinson’s Research and funding partners, including 4D Pharma, Abbvie, AcureX, Allergan, Amathus Therapeutics, Aligning Science Across Parkinson’s, AskBio, Avid Radiopharmaceuticals, BIAL, BioArctic, Biogen, Biohaven, BioLegend, BlueRock Therapeutics, Bristol-Myers Squibb, Calico Labs, Capsida Biotherapeutics, Celgene, Cerevel Therapeutics, Coave Therapeutics, DaCapo Brainscience, Denali, Edmond J. Safra Foundation, Eli Lilly, Gain Therapeutics, GE HealthCare, Genentech, GSK, Golub Capital, Handl Therapeutics, Insitro, Jazz Pharmaceuticals, Johnson & Johnson Innovative Medicine, Lundbeck, Merck, Meso Scale Discovery, Mission Therapeutics, Neurocrine Biosciences, Neuron23, Neuropore, Pfizer, Piramal, Prevail Therapeutics, Roche, Sanofi, Servier, Sun Pharma Advanced Research Company, Takeda, Teva, UCB, Vanqua Bio, Verily, Voyager Therapeutics, the Weston Family Foundation and Yumanity Therapeutics. We would like to thank all of the subjects who donated their time and biological samples to PPMI. We also thank Peter Heutink for initial support and discussions. This work was funded in part by The Michael J. Fox Foundation for Parkinson’s Research, MIT MISTI Global Seed Fund and a DFG grant (Project number 513977564 to V.B.). V.B. is also supported by a Career Development Fellowship at DZNE Tuebingen. BioRender was used to prepare Figure 1A.

## Supplementary table legends

**Table S1**: Sample overview with meta-data

**Table S2**: Genotype check for sample concordance and identity

**Table S3**: Number of cells across broad cell types and mutation group

**Table S4**: Known and identified cluster markers.

**Table S5**: MAGMA scRNA-seq cell-type enrichment results.

**Table S6**: Number of cells across cell types and mutation group with speckle analysis

**Table S7**: Differentially expressed genes between LRRK2 and HC in each cell type

**Table S8**: Enriched gene ontology terms for DEGs for all subclusters from EnrichR

**Table S9**: Cell-cell communication information flow statistics

**Table S10**: Trajectory-based genes associated with pseudotime and differential expression

**Table S11**: Description of antibodies for ICC and FACS

